# Cancer LncRNA Census 2 (CLC2): an enhanced resource reveals clinical features of cancer lncRNAs

**DOI:** 10.1101/2020.04.28.066225

**Authors:** Adrienne Vancura, Andrés Lanzós, Núria Bosch, Mònica Torres, Alejandro Hionides Gutierrez, Simon Haefliger, Rory Johnson

## Abstract

Long noncoding RNAs play key roles in cancer and are at the vanguard of precision therapeutic development. These efforts depend on large and high-confidence collections of cancer lncRNAs. Here we present the Cancer LncRNA Census 2 (CLC2): at 492 cancer lncRNAs, it is 4-fold greater than its predecessor, without compromising on strict criteria of confident functional / genetic roles and inclusion in the GENCODE annotation scheme. This increase was enabled by leveraging high-throughput transposon insertional mutagenesis (TIM) screening data, yielding 95 novel cancer lncRNAs. CLC2 makes a valuable addition to existing collections: it is amongst the largest, holds the greatest number of unique genes, and carries functional labels (oncogene / tumour suppressor). Analysis of this dataset reveals that cancer lncRNAs are impacted by germline variants, somatic mutations, and changes in expression consistent with inferred disease functions. Furthermore, we show how clinical / genomic features can be used to vet prospective gene sets from high-throughput sources. The combination of size and quality makes CLC2 a foundation for precision medicine, demonstrating cancer lncRNAs’ evolutionary and clinical significance.

## Introduction

Tumours arise and grow via genetic and non-genetic changes that give rise to widespread alterations gene expression programmes (1–3). The numerous dysregulated genes may encode classical protein-coding mRNAs or non-protein coding RNAs, but it is likely that just a subset of these actually functionally contribute to pathogenic cellular hallmarks. The identification of such functional cancer genes is critical for the development of targeted cancer therapies, as well as emerging methods to identify additional cancer genes. For protein-coding genes (pc-genes), datasets such as the Cancer Gene Census (CGC) collect and organise comprehensive gene collections according to defined criteria, and has proven invaluable for scientific research and drug discovery (4).

The past decade has witnessed the discovery of numerous non-protein-coding RNA genes in mammalian cells (5, 6). The most numerous but poorly understood produce long noncoding RNAs (lncRNAs), defined as transcripts >200 nt in length with no detectable protein-coding potential (7). Although their molecular mechanisms are highly diverse, many lncRNAs have been shown to interact with other RNA molecules, proteins and DNA by structural and sequence-specific interactions (8, 9). Most lncRNAs are clade- and species-specific, but a subset display deeper evolutionary conservation in their gene structure (10) and a handful have been demonstrated to have functions that were conserved across millions of years of evolution (10, 11). The numbers of known lncRNA genes in human have grown rapidly, and present catalogues range from 18,000 to ~100,000 (12), however just a tiny fraction have been functionally characterized (13–16). As lncRNAs likely represent a huge yet poorly understood component of cellular networks, understanding the clinical and therapeutic significance of these numerous novel genes is a key contemporary challenge.

LncRNAs have been implicated in molecular processes governing tumorigenesis (17). LncRNAs may promote or oppose cancer hallmarks (18). This fact, coupled to the emergence of potent *in vivo* inhibitors in the form of antisense oligonucleotides (ASOs) (19), has given rise to serious interest in lncRNAs as drug targets in cancer by both academia and pharma (17, 20–22).

Initially, cancer lncRNAs were discovered by classical functional genomics workflows employing microarray or RNA-seq expression profiling (23, 24). More recently, CRISPR-based functional screening (25) and bioinformatic predictions (26–28) have also emerged as powerful tools for novel cancer gene discovery. To assess their accuracy, these approaches require accurate benchmarks in the form of curated databases of known cancer lncRNAs.

Any discussion of lncRNAs and cancer requires careful terminology. Experimental evidence suggest that for some lncRNAs, it is a DNA element within the gene, in addition to or instead of the RNA transcript, which mediates downstream gene regulation (29–31). This introduces the need for meticulous assessment of the basis of each lncRNA gene’s functionality. Furthermore, it has been shown that lncRNAs can exert strong phenotypic effects in one cell background, but none in another (32). In the context of tumours, this means that amongst the large numbers of differentially expressed lncRNAs (24), just a fraction are likely to functionally contribute to a relevant cellular phenotype or cancer hallmark (20, 33–36). Such genes, termed here “functional cancer lncRNAs”, are the focus of this study. Remaining changing genes are non-functional “bystanders”, which are largely irrelevant in understanding or inhibiting the molecular processes causing cancer and highlight the importance of not assessing functionality evidence simply by expressional changes.

There are a number of excellent databases of cancer-associated lncRNAs: lncRNADisease (37), CRlncRNA (38), EVLncRNAs (39) and Lnc2Cancer (40). These principally employ labour-intensive manual curation, and rely extensively on differential expression to identify candidates. On the other hand, these databases have not yet taken advantage of recent high-confidence sources of functional cancer lncRNAs, such as high-throughput functional screens (25, 41). For these reasons, existing annotations likely contain unknown numbers of bystander lncRNAs, while omitting large numbers of *bona fide* functional cancer lncRNAs. Thus, studies requiring high-confidence gene sets, including benchmarking or drug discovery, call for a database focussed exclusively on functional cancer lncRNAs.

Here we address this need through the creation of the Cancer LncRNA Census 2 (CLC2). It not only extends our previous CLC dataset by several fold (42), but more importantly, CLC2 takes a major step forward methodologically, by implementing an automated curation component that utilises functional evolutionary conservation for the first time. Using this data, we present a comprehensive analysis of the genomic and clinical features of cancer lncRNAs.

## Results

### Integrative, semi-automated cataloguing of cancer lncRNAs

We sought to develop an improved map of lncRNAs with functional roles in either promoting or opposing cancer hallmarks or tumorigenesis. Such a map should prioritise lncRNAs with genuine causative roles, and exclude false-positive “bystanders”: genes whose expression changes but play no functional role.

We began with conventional manual curation of lncRNAs from the scientific literature, covering the period from January 2017 (directly after the end of the first CLC (42)) to the end of December 2018. We continued to use stringent criteria for defining cancer lncRNAs: genes must be annotated in GENCODE (here version 28), and cancer function must be demonstrated either by functional *in vitro* or *in vivo* experiments, or germline or somatic mutational evidence (see Methods) (Figure 1A). Altogether we collected 253 novel lncRNAs in this way, which added to the original CLC amounts to 375 lncRNAs, hereafter denoted as “literature lncRNAs” (Figure 1A).

**Figure 1:**
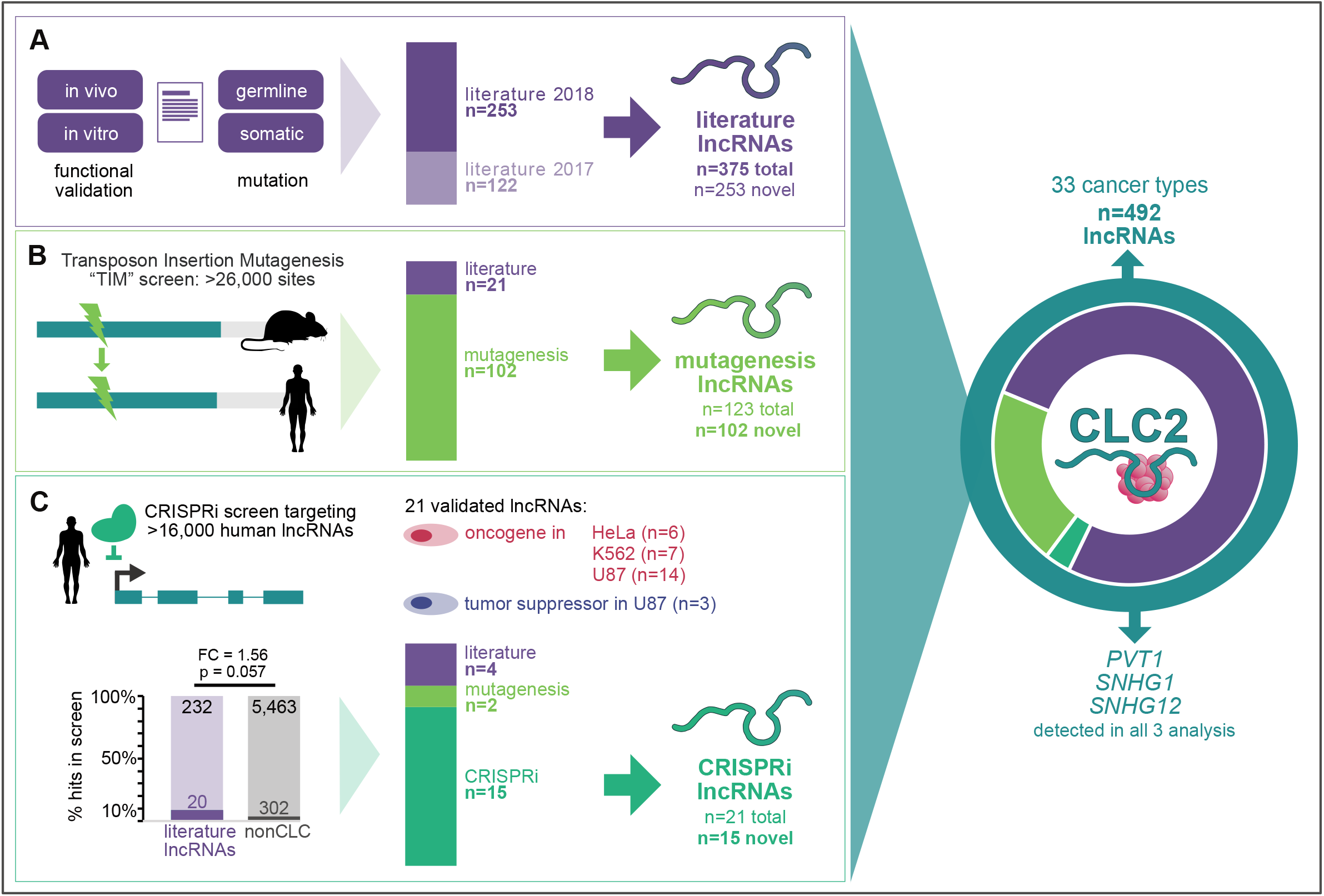
Functional cancer lncRNAs from three sources are integrated in the CLC2. **A)** Literature curation with four criteria are used to define “literature lncRNAs”. **B)** Transposon insertion mutagenesis screens identify “mutagenesis lncRNAs”. **C)** Validated hits from CRISPRi proliferation screens are denoted “CRISPRi lncRNAs”. Statistical significance calculated by one-sided Fisher’s test.

We recently showed that some literature-curated lncRNAs were also targeted by previously-overlooked mutations in published transposon insertion mutagenesis (TIM) screens (42). We hypothesised that this insight could be extended to identify novel functional cancer lncRNAs. Thus we developed a pipeline to automatically identify human lncRNAs by orthology to a collection of TIM hits in mouse (41). In this way 123 lncRNAs were detected, of which 102 were not already in the literature set. These were added to the CLC2, henceforth denoted as “mutagenesis lncRNAs” (Figure 1B). This analysis is discussed in more detail in the next section.

Pooled functional screens based on CRISPR-Cas9 loss-of-function have recently emerged as a powerful means of identifying function cancer lncRNAs (25). However there has been relatively little validation of the hits from such screens, and it is possible that they contain substantial false positives (43, 44). Amongst the few datasets presently available, the most comprehensive comes from a CRISPR-inhibition (CRISPRi) screen of ~16,000 lncRNAs in seven human cell lines, with proliferation as a readout (45). Of the 499 hits identified, 322 are annotated by GENCODE and hence could potentially be included in CLC2. These are moderately enriched for known cancer lncRNAs from the literature search (Figure 1C). That study independently validated 21 GENCODE-annotated hits, of which four (19%) were already mentioned in the literature, and two (10%) were detected by TIM above. Given the uncertainty over the true-positive rates of unvalidated screen hits, we opted for a conservative approach and included the remaining 15 novel and independently-validated lncRNAs from this study (“CRISPRi lncRNAs”) (Figure 1C).

Altogether, the resulting CLC2 set comprises 492 unique lncRNA genes, representing a 4.0-fold increase over its predecessor. The entire CLC2 dataset is available in Supplementary Table 1 and 2. Importantly, the dataset is fully annotated with evidence information, affording users complete control over the particular subsets of lncRNAs (literature, mutagenesis, CRISPRi) that they wish to include in their analyses.

### Automated annotation of human cancer lncRNAs via functional conservation

We recently showed that transposon insertional mutagenesis (TIM) screens identify cancer lncRNAs in mouse (42, 46), and that some of these overlapped previously-known human cancer lncRNAs (Figure 2A). TIM screens identify “common insertion sites” (CIS), where multiple transposon insertions at a particular genomic location have given rise to a tumour, thereby implicating the underlying gene as an oncogene or tumour suppressor.

**Figure 2:**
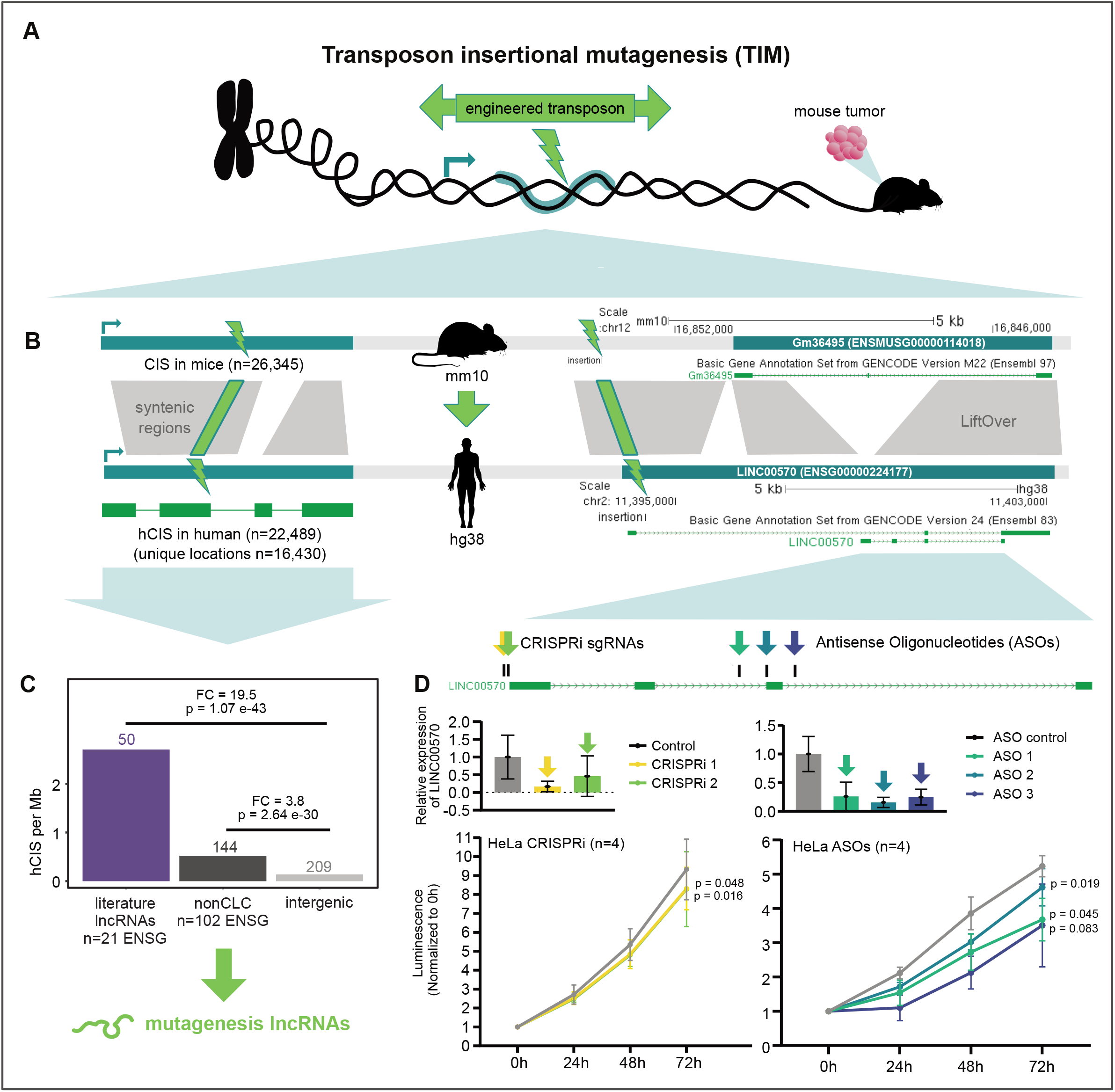
The CLIO-TIM pipeline identifies human cancer lncRNAs via functional evolutionary conservation. **A)** Overview of transposon insertional mutagenesis (TIM) method for identifying functional cancer genes. Engineered transposons carry bidirectional cassettes capable of either blocking or upregulating gene transcription, depending on orientation. Transposons are introduced into a population of cells, where they integrate at random genomic sites. The cells are injected into a mouse. In some cells, transposons will land in and perturb expression of a cancer gene (either tumour suppressor or oncogene), giving rise to a tumour. DNA of tumour cells is sequenced to identify the exact location of the transposon insertion. Clusters of such insertions are termed Common Insertion Sites (CIS). **B)** (Left) Schematic of the CLIO-TIM pipeline used here to identify human cancer genes using mouse CIS. (Right) An example of a CLIO-TIM predicted cancer lncRNA. **C)** The density of hCIS sites, normalised by gene length, in indicated classes of lncRNAs. Statistical significance calculated by one-sided Fisher’s test. **D)** Upper panels: Expression of *LINC00570* RNA in response to inhibition by CRISPRi (left) or ASOs (right). Lower panels: Measured populations of the same cells over time. Statistical significance calculated by Student’s *t*-test.

We here extend this strategy to identify new functional cancer lncRNAs, by developing a new pipeline called CLIO-TIM (cancer lncRNA identification by orthology to TIM). Briefly, CLIO-TIM uses chain alignments (47) to map mouse CIS to orthologous regions of the human genome, and then identifies the most likely gene target (see Methods) (Figure 2B) (SUPP FIG 1B). Available CIS maps are based on a variety of identification methods, resulting in CIS with a range of sizes, from 1 bp upwards. We opted to remove our previously conservative size criterion (CIS = 1 bp), to now consider elements of any size resulting in 26,345 CIS (compared to 2,806 previously (42)) (SUPP FIG 1A). This yields a 3-fold increase in sensitivity for true-positive CGC genes (72% compared to 26.4% previously (42)) (SUPP FIG 1D).

Based on this expanded dataset, CLIO-TIM identified 16,430 orthologous regions in human (hCIS) (Figure 2B) (SUPP FIG 1A). Altogether, 123 lncRNAs and 9,295 pc-genes were identified as potential cancer genes. An example is the human-mouse orthologous lncRNA locus shown in Figure 2B, comprising *Gm36495* in mouse and *LINC00570* in human. A CIS lies upstream of the mouse gene’s TSS, mapping to the first intron of the human orthologue. *LINC00570* is an alternative identifier for ncRNA-a5 *cis*-acting lncRNA identified by Orom et al. (48), that has not previously been associated with cancer or cell growth.

We expected that hCIS regions are enriched in known cancer genes. Consistent with this, the 698 pc-genes from the COSMIC Cancer Gene Census (CGC) (4) (red in SUPP FIG 1D) are 155-fold enriched with hCIS over intergenic regions (light grey). Turning to lncRNAs, the 375 literature lncRNAs are 19.5-fold enriched, supporting their disease relevance (Figure 2C). Thus, CLIO-TIM predictions are enriched in genuine protein-coding and lncRNA functional cancer genes. Supporting its accuracy, the overall numbers of genes implicated by CLIO-TIM agree with independent analysis in the CCGD database (SUPP FIG 1C).

An additional 209 hCIS fall in intergenic regions that are neither part of pc-genes or lncRNAs, leading us to ask whether some may affect lncRNAs that are not annotated by GENCODE (Figure 2C). To test this, we utilised the large set of cancer-associated lncRNAs from miTranscriptome (24). 186 hCIS intersect 2167 miTranscriptome genes, making these potentially novel non-annotated transcripts involved in cancer. Nevertheless, simulations indicated that this rate of overlap was no greater than expected by random chance (see Methods), making it unlikely that substantial numbers of undiscovered cancer lncRNAs remain to be discovered in intergenic regions, at least with the datasets used here (SUPP FIG 1E).

In addition to known cancer lncRNAs, CLIO-TIM identifies 102 lncRNAs not previously linked to cancer (FIG 2C, dark grey) with a 3.8-fold enrichment of insertions over intergenic genome. As will be shown below, these lncRNAs bear clinical and genomic features of functional cancer genes, and hence we decided to include them in CLC2. It should be noted, however, that these “mutagenesis” lncRNAs are labelled and hence may be removed by end users, as desired.

To experimentally test the principal that human orthologues of mouse cancer genes have a conserved function, we selected *LINC00570*, identified by CLIO-TIM but never previously been linked to cancer or cell proliferation. We asked whether *LINC00570* promotes cell growth in transformed cells. We used RNA-sequencing data to search for cell models where LINC00570 is expressed, and identified robust expression in cervical carcinoma HeLa cells (SUPP FIG 2A) and to a lesser extent in HCT116 colon carcinoma cells (SUPP FIG 2A). We designed three distinct antisense oligonucleotides (ASOs) targeting the *LINC00570* intron 2 and 3 and exon 3 of the short isoform (intronic targeting ASOs are known to have degradation efficiency comparable to exonic ones (49, 50)). Transfection of these ASOs led to strong and reproducible decreases in steady state RNA levels in HeLa cells (Figure 2C). This resulted in significant decreases in cell proliferation rates (Figure 2D, SUPP FIG 2B). We observed a similar effect through CRISPRi-mediated inhibition of gene transcription by two independent guide RNAs in HeLa (Figure 2D), and with the same ASOs in HCT116 cells (SUPP FIG 2C and D). Therefore, *LINC00570* predicted by CLIO-TIM pipeline promotes growth of human cancer cells, and is likely to have a deeply evolutionarily-conserved tumorigenic activity.

### Enhanced cancer lncRNA catalogue integrating manual annotation, CRISPR screens and functional conservation

We next tallied the distinct lncRNAs in CLC3 and compared them with existing cancer lncRNA databases. Figure 3A shows a breakdown of the composition of CLC2 in terms of source, gene function and evidence strength. Where possible, the genes are given a functional annotation, oncogene (og) or tumour suppressor (ts), according to evidence for promoting or opposing cancer hallmarks. Oncogenes (n=275) quite considerably outnumber tumour suppressors (n=95), although it is not clear whether this reflects genuine biology or an ascertainment bias relating to scientific interest or technical issues. Smaller sets of lncRNAs are associated with both functions, or have no functional information (those from TIM screens where the functions of hits are ambiguous).

**Figure 3:**
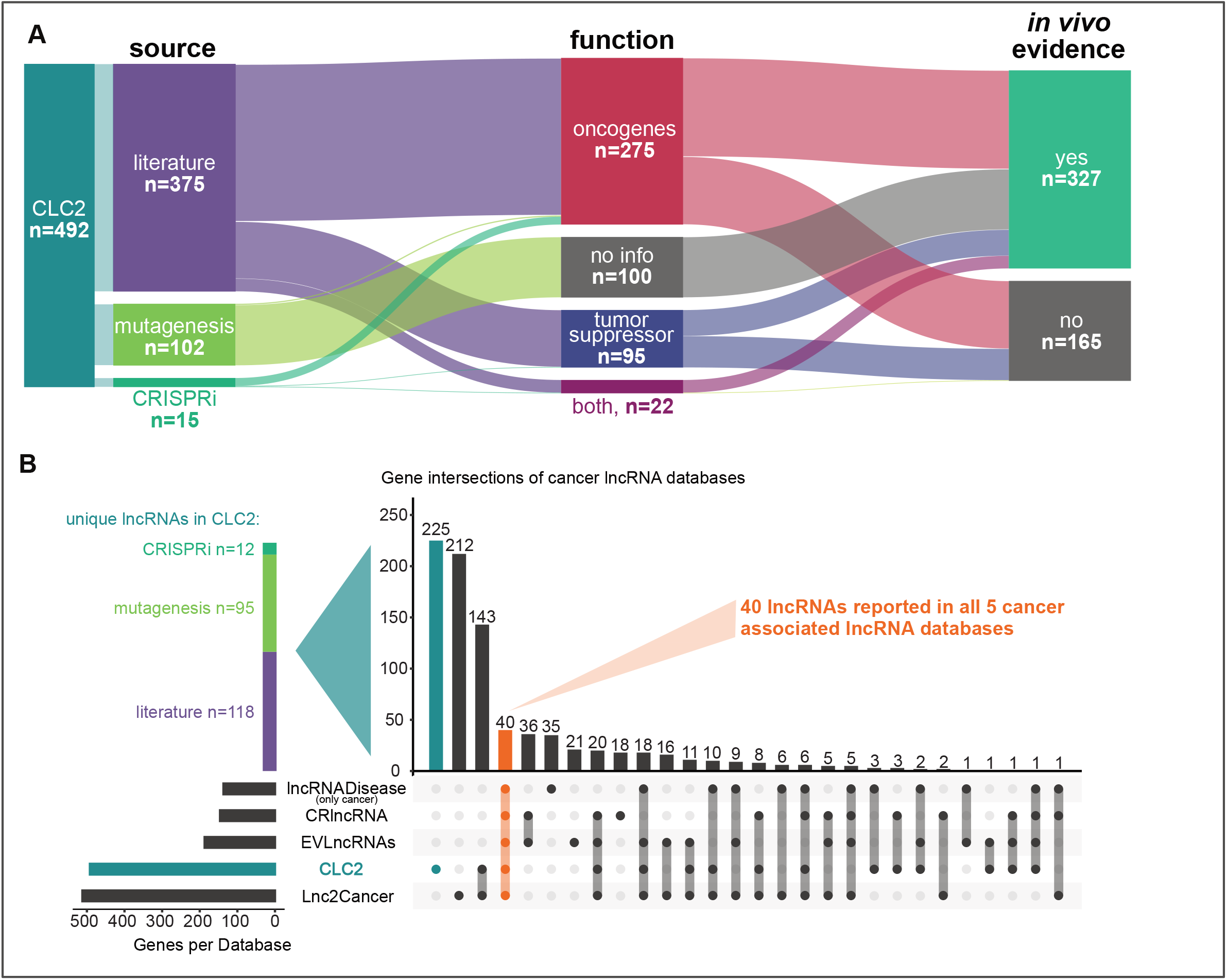
An overview of the CLC2 database and comparison with other lncRNA databases. **A)** The CLC2 database broken down by source, function and evidence type. **B)** Comparison of CLC2 to other leading cancer lncRNA databases.

In terms of the quality of evidence sources, CLC2 represents a considerable improvement over the original CLC. The fraction of lncRNAs with high quality *in vivo* evidence (defined as functional validation in mouse models or mutagenesis analysis) now represent 66% compared to 24% previously (Figure 3A, SUPP FIG 3B). In total, the updated CLC2 comprises 33 cancer types (vs 29) and more lncRNAs are reported for every cancer subtype (SUPP FIG 3A).

We were curious how much novelty the CLC2 gene set brought to the known universe of cancer lncRNAs, as estimated from respected and longstanding cancer lncRNA collections (Figure 3B). Considering only GENCODE-annotated genes, CLC2 with 492 is second only to Lnc2Cancer (n= 512) in terms of size (40). However, Lnc2Cancer uses looser inclusion criteria, including lncRNAs that are differentially expressed in tumours without additional functional evidence. The three remaining databases are smaller (<200 genes). Importantly, CLC2 holds the greatest number of unique genes, i.e. those that are not found in other databases (n=225). These contain 118 literature-annotated cases, and also 95 novel mutagenesis lncRNAs. Just 40 lncRNAs are common to all five databases (37–40). In summary, CLC2 achieves large size without compromising on confidence, while also including numerous new cancer lncRNAs for the first time.

### Unique genomic properties of CLC2 lncRNAs

Cancer genes, both protein-coding and not, display elevated characteristics of essentiality and clinical importance compared to other genes (4, 18, 51, 52). In order to confirm their quality as a resource, we next asked whether CLC2 lncRNAs, and the mutagenesis subset, display features expected for cancer genes.

In the following analyses, we compared gene features of selected lncRNAs to all other lncRNAs. Comparison of gene sets can often be confounded by covariates such as gene length or gene expression, therefore where appropriate we used control gene sets that were matched to CLC2 by expression (denoted “nonCLCmatched”) (SUPP FIG 4A) and reported findings correcting for gene length (SUPP FIG 4B).

Evolutionary conservation and steady-state expression are widely-used proxies for gene function (53–55). Using the LnCompare tool (56), we find that the promoters and exons of CLC2 genes display elevated evolutionary conservation in mammalian and vertebrate phylogeny (Figure 4A) and elevated expression in cancer cell lines (Figure 4B). Strikingly we observe a similar effect when considering the mutagenesis lncRNAs alone: their promoters are significantly more conserved than expected by chance, and their expression is an order of magnitude higher than other lncRNAs (Figure 4C and D).

**Figure 4:**
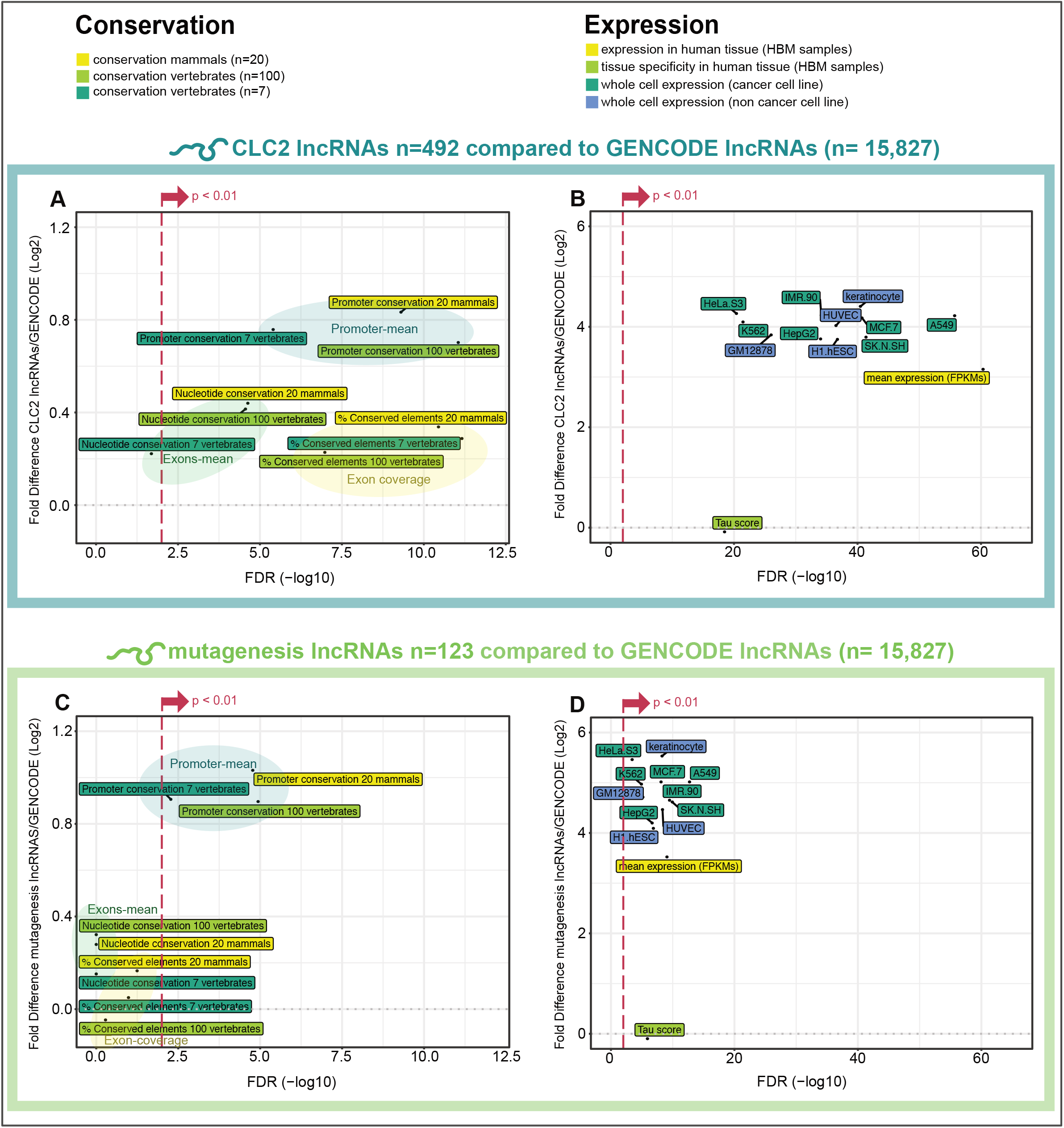
Features of functionality in CLC2 and mutagenesis lncRNAs. In each panel, two gene sets are compared: the test set (either all CLC2 genes, or mutagenesis genes alone), and the set of all other lncRNAs (GENCODE v24). Y-axis: Log2 fold difference between the means of gene sets. X-axis: false-discovery rate adjusted significance, calculated by Wilcoxon test. **A)** Evolutionary conservation for all CLC2, calculated by PhastCons. **B)** Expression of all CLC2 in cell lines. **C)** Evolutionary conservation for mutagenesis lncRNAs, calculated by PhastCons. **D)** Expression of mutagenesis lncRNAs in cell lines. For (A) and (C), “Promoter mean” and “Exon mean” indicate mean PhastCons scores (7-vertebrate alignment) for those features, while “Exon-coverage” indicates percent coverage by PhastCons elements. Promoters are defined as a window of 200 nt centered on the transcription start site.

Further, we found that CLC2 lncRNAs are enriched in repetitive elements (SUPP FIG 5A) and are more likely to house a small RNA gene, possibly indicating that some act as precursor transcripts (SUPP FIG 5B). CLC2 lncRNAs also have non-random distributions of gene biotypes, being depleted for intergenic class and enriched in divergent orientation to other genes (SUPP FIG 5C).

In summary, CLC2 lncRNAs are significantly more conserved and more expressed than expected by chance, pointing to biological function. Mutagenesis lncRNAs discovered by the CLIO-TIM also carry these features, supports their designation as functional cancer lncRNAs.

### CLC2 lncRNAs display consistent tumour expression changes and prognostic properties

Although gene expression was not a criterion for inclusion, we would expect that CLC2 lncRNAs’ expression will be altered in tumours. Furthermore, one might expect that the nature of this alteration should vary with disease function: oncogenes overexpressed, and tumour suppressors downregulated.

To test this, we analysed TCGA RNA-sequencing (RNA-seq) data from 686 individual tumours with matched healthy tissue (total n=1,372 analysed samples) in 20 different cancer types (SUPP FIG 6A and B), and classified every gene as either differentially expressed (in at least one cancer subtype, with log2 Fold Change >1 and FDR <0.05) or not. We found that CLC2 lncRNAs are 3.4-fold more likely to be differentially expressed compared to expression-matched lncRNAs (Figure 5A). LncRNAs from each individual evidence source (literature, mutagenesis, CRISPRi) behaved similarly, again supporting their inclusion. Similar effects were found for pc-genes (SUPP FIG 7A).

**Figure 5:**
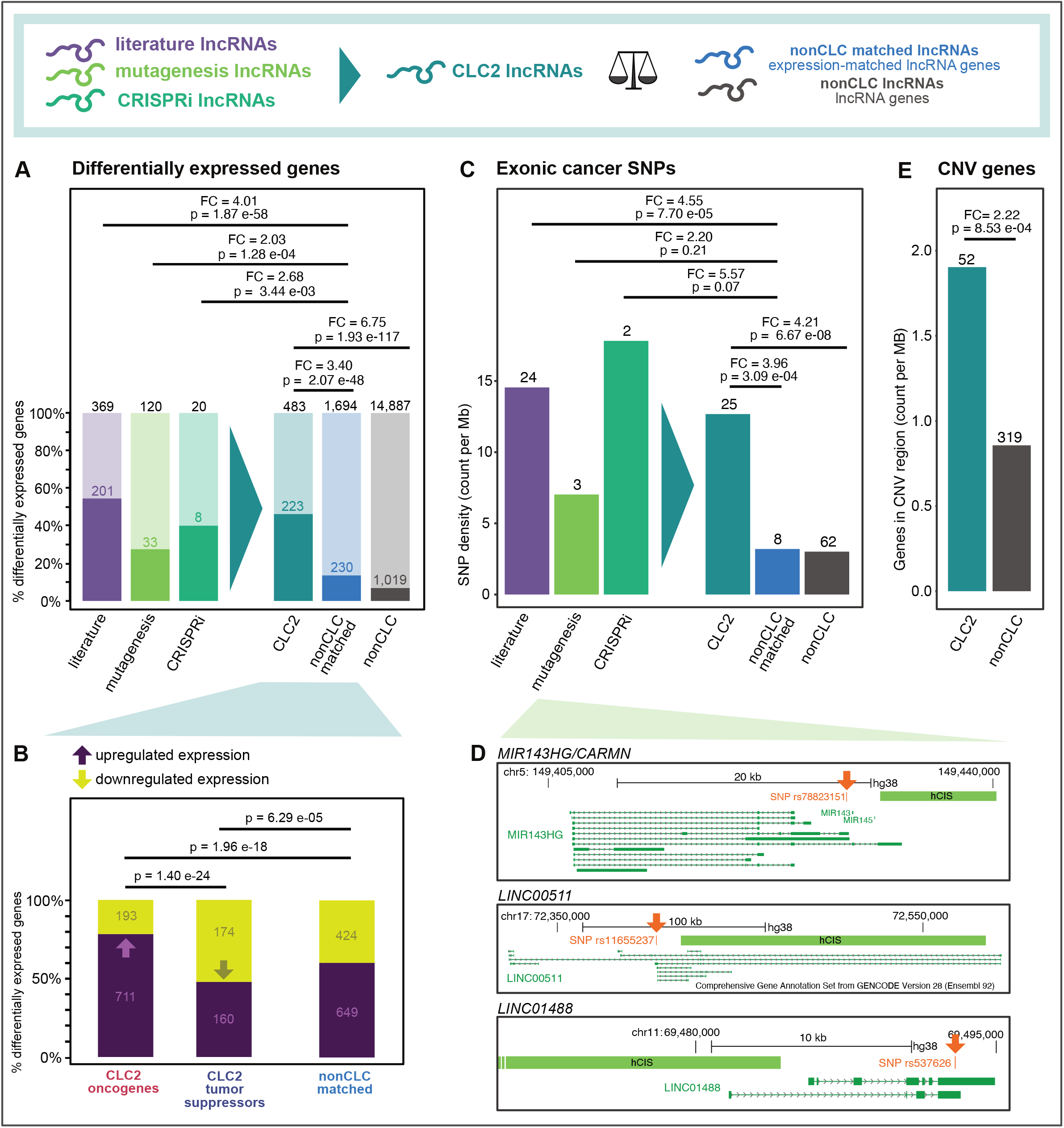
Clinical features of CLC2 lncRNAs. **A)** The percent of indicated genes that are significantly differentially expressed in at least one tumour type from the TCGA. Statistical significance calculated by one-sided Fisher’s test. **B)** Here, only differentially expressed genes from (A) are considered. LncRNAs with both tumour suppressor and oncogene labels are excluded. Remaining lncRNAs are divided by those that are up- or down-regulated (positive or negative fold change). Statistical significance calculated by one-sided Fisher’s test. **C)** The density of germline cancer-associated SNPs is displayed. Only SNPs falling in gene exons are counted, and are normalised to the total length of those exons. Statistical significance calculated by one-sided Fisher’s test. **D)** Examples of mutagenesis lncRNAs with an exonic cancer SNP. **E)** Length-normalised overlap rate of copy number variants (CNVs) in lncRNA gene span. Statistical significance calculated by one-sided Fisher’s test.

Next, we asked whether the direction of expression change corresponds to gene function. Indeed, oncogenes are enriched for overexpressed genes, whereas tumour suppressors are enriched for down-regulated genes, supporting the functional labelling scheme (Figure 5B).

Cancer genes’ expression is often prognostic for patient survival. By correlating expression to patient survival, we found that the expression of 392 CLC2 lncRNAs correlated to patient survival in at least one cancer type (SUPP FIG 7C). When analysing the most significant correlation of each CLC2 lncRNA compared to expression-matched nonCLC lncRNAs, we find a weak but significant enrichment (SUPP FIG 7C), suggesting that CLC2 lncRNAs can be prognostic for patient survival.

In summary, gene expression characteristics of CLC2 genes, and subsets from different evidence sources, support their functional labels as oncogenes and tumour suppressors and is more broadly consistent with their important roles in tumorigenesis.

### CLC2 lncRNAs are enriched with cancer genetic mutations

Cancer genes are characterized by a range of germline and somatic mutations that lead to gain- or loss-of-function. It follows that cancer lncRNAs should be enriched with germline single nucleotide polymorphisms that have been linked to cancer predisposition (57). We obtained 5,331 germline cancer-associated single nucleotide polymorphisms (SNPs) from genome-wide association studies (GWAS) (58) and mapped them to lncRNA and pc-gene exons, calculating a density score that normalises for exon length (SUPP FIG 4B). As expected, exons of known cancer pc-genes are >2-fold enriched in cancer SNPs (SUPP FIG 7B). When performing the same analysis with CLC2 lncRNAs, one observes an even more pronounced enrichment of 4.0-fold when comparing to expression-matched nonCLC lncRNAs (Figure 5C). Once again, the lncRNAs from each evidence source individually show enrichment for cancer SNPs >2-fold (Figure 5C). Three mutagenesis lncRNAs, namely *miR143HG/CARMN*, *LINC00511* and *LINC01488*, carry an exonic cancer SNP (Figure 5D).

Cancer genes are also frequently the subject of large-scale somatic mutations, or copy number variants (CNVs). Using a collection of CNV data from LncVar (59), we calculated the gene-span length-normalized coverage of lncRNAs by CNVs. CLC2 lncRNAs are enriched for CNVs compared to all lncRNAs (Figure 5E).

All information of the lncRNAs in the CLC2 with the corresponding cancer function, evidence level, analysis method and cancer types can be found in the Supplementary Table 1. The Supplementary Table 2 can be used to filter lncRNAs based on their reported cancer associated functionalities.

In summary, CLC2 lncRNAs and their subsets display germline and somatic mutational patterns consistent with known oncogenes and tumour suppressors.

## Discussion

We have presented the Cancer LncRNA Census 2, an expanded collection of lncRNAs with functional roles in cancer. CLC2 is distinguished from other resources by several key features. All its constituent lncRNAs have strong evidence for functional cancer roles (and not merely differential expression), providing for lowest possible false positive rates. All CLC2 lncRNAs are included in the gold-standard GENCODE annotation, permitting smooth interoperability with almost all public genomics projects and resources (12). The majority of CLC2 entries are accompanied by functional labels (oncogene / tumour suppressor), enabling one to link function to other observable features. Finally, we utilise transposon insertional mutagenesis (TIM) datasets for the first time to discover 102 “mutagenesis” lncRNAs, of which 95 are completely novel. In spite of strict inclusion criteria, CLC2 is amongst the largest available cancer lncRNA collections. Most striking, is that it contains the greatest number of “unique” lncRNAs, not found in other resources. Overall, CLC2 makes a valuable addition to the present landscape of cancer lncRNA resources.

A key novelty of CLC2 is its use of automated gene curation based on functional evolutionary conservation, as inferred from TIM. This responds to the challenge from the rapid growth of scientific literature, which makes manual curation increasingly impractical. Other high-throughput / automated methods like CRISPR pooled screening, text mining and machine learning will also be important, although it will be necessary to vet the quality of such predictions prior to inclusion. Here we showed one way approach for this, by assessing the TIM gene set across a range of genomic and clinical features. The fact that the “mutagenesis” lncRNA set display rates of (i) nucleotide conservation, (ii) expression, (iii) tumour differential expression, (iv) germline cancer polymorphisms and (v) tumour mutations similar to that of gold-standard literature curated lncRNAs, coupled to thorough experimental validation of one novel prediction (*LINC00570*), is powerful support for TIM and functional evolutionary conservation as means for new cancer lncRNA discovery.

It might be argued that hits from TIM sites could be false positives that act via DNA elements (for example, enhancers) that, by coincidence, overlap a non-functional lncRNA. While certainly likely to occur in some cases, it would nevertheless appear unlikely to explain the majority, in light of the features listed above, plus the observation that TIM sites are highly enriched in independently-validated literature-curated lncRNAs (which act via RNA) including *NEAT1*, *LINC-PINT* and *PVT1* (42). In spite of this, we recognise that some colleagues may ascribe lower confidence to novel “mutagenesis” lncRNAs in CLC2. For this reason, the CLC2 data table is organised to facilitate filtering by source, enabling users to extract only the 375 literature-supported cases, or indeed any other subset based on source, evidence or function as desired.

Apart from its usefulness as a resource, this study has enabled some important conceptual insights. Firstly, we have replicated our previous finding that cancer lncRNAs are distinguished by signatures of functionality, as inferred from evolutionary nucleotide conservation and expression. These features were originally linked to protein-coding cancer genes (51, 52), but are also utilised as markers for lncRNA functionality (42, 60). Moreover, we extended this approach to clinical features, by showing that curated cancer lncRNAs are dramatically more likely to be differentially expressed in tumours, suffer copy number alteration, or carry a germline predisposition SNP. In the latter case, this rate even exceeds cancer driver protein-coding genes. We also could demonstrate that changes in gene expression in tumours are linked to function: oncogenes tend to be overexpressed, while tumour-suppressors tend to be repressed. Finally, the demonstration that cancer lncRNAs can be predicted on the basis of orthology to a TIM hit in mouse, lends powerful support to the notion that there is widespread functional evolutionary conservation of lncRNAs in networks related to cell growth and transformation.

*LINC00570* is a new functional cancer lncRNA predicted by CLIO-TIM. The gene was previously discovered by Orom and colleagues, as a *cis*-activating enhancer-like RNA named *ncRNA-a5* (48). That and a subsequent study showed that perturbation by siRNA transfection affects the expression of the nearby pc-gene *ROCK2* in HeLa. However, these studies did not investigate the effect on cell proliferation. We here show by means of two independent perturbations, that *LINC00570* promotes proliferation of HeLa and HCT116 cells. These findings make *LINC00570* a potential therapeutic target for follow up.

Intriguingly, amongst the novel mutagenesis lncRNAs identified by CLIO-TIM are genes previously linked to other diseases. *miR143HG/CARMEN1* (*CARMN*) was shown to regulate cardiac specification and differentiation in mouse and human hearts (61). In addition to being a TIM target, CARMEN1 also contains a germline cancer SNP correlating to the risk of developing lung cancer (62), adding further weight to the notion that it also plays a role in oncogenesis. Similarly, *DGCR5*, is located in the DiGeorge critical locus and has been linked to neurodevelopment and neurodegeneration (63), and was recently implicated as a tumour suppressor in prostate cancer (64). These results raise the possibility that developmental lncRNAs can also play roles in cancer.

In summary, CLC2 establishes a new benchmark for cancer lncRNA resources. We hope this dataset will enable a wide range of studies, from bioinformatic identification of new disease genes, to developing a new generation of cancer therapeutics with anti-lncRNA ASOs (65).

## Material and Methods

### Gene curation

If not stated otherwise, GENCODE v28 gene IDs (gencode.v28.annotation.gtf) were used.

#### Literature search

PubMed was searched for publications linking lncRNA and cancer using keywords: long noncoding RNA cancer, lncRNA cancer. Additional inclusion criteria consisted of GENCODE annotation, reported cancer subtype and cancer functionality (oncogene/tumour suppressor). The manual curation and assigning evidence levels to each lncRNA was performed exactly as previously (42) and included reports until December 2018.

#### CLIO-TIM

From the CCGD website (http://ccgd-starrlab.oit.umn.edu/about.php, May 2018 (41)) a table with all CIS elements was downloaded. These mouse genomic regions (mm10) were converted to homologous regions in the human genome assembly hg38 using the LiftOver tool (https://genome.ucsc.edu/cgi-bin/hgLiftOver). Settings: original Genome was Mouse GRCm38/mm10 to New Genome Human GRCh38/hg38, minMatch was 0.1 and minBlocks 0.1. For insertion sites intersecting several lncRNA genes, all the genes were reported. IntersectBed from bedtools was used to align human insertion sites to GENCODE IDs by intersecting at least 1nt and assigned to protein-coding or lncRNA gene families. Insertion sites aligning to protein-coding and lncRNA genes were always assigned to protein-coding genes. If insertion sites overlap multiple ENSGs, all genes are reported. Insertion sites not aligning to protein-coding or lncRNAs genes were added to the intergenic region.

CCGD human Entrez gene results were converted to GENCODE IDs using the “Entrez gene ids” Metadata file from https://www.gencodegenes.org/human/ to compare CLIO-TIM results with CCGD results for each gene set.

#### MiTranscriptome data for evaluating intergenic insertion sites

The cancer associated MiTranscriptome IDs (24) previously used in Bergada et al. (66) were intersected with intergenic insertion sites using IntersectBed. With ShuffleBed the intergenic insertions were randomly shuffled 1000x and assigned to MiTranscriptome IDs.

#### CRISPRi

We used the Supplementary Table 1 from the 2017 Liu et al. paper (45) to extract ENST IDs and gene names which are then converted to GENCODE IDs to match each guide (LH identifier in the screen). From Supplementary Table S4 from the 2017 Liu et al. paper (Liu_et_al_aah7111-TableS4) (45) we extracted genes with “hit” (validated as a hit in the screen), “LH” (unique identifiers correlating to a gene in the screen) and “lncRNA” (referring to a lncRNA gene and to exclude lncRNA hits close to a protein-coding gene (“Neighbor hit”)) resulting in 499 hits. Of these, 322 hits contain a GENCODE IDs and were used for enrichment analysis, tested by one-sided Fisher’s test.

We included n=21 CRISPRi genes to the CLC2 from the Supplementary Figure 8A from the 2017 Liu et al. paper (45), the tested cancer cell line and the effect of the CRISPRi on the growth phenotype (either promoting (tumor suppressor) or inhibiting (oncogene)) of each lncRNA was reported.

#### Cancer gene sets

For downstream analysis protein-coding (pc) genes (GENCODE IDs) are grouped in cancer-associated pc-genes (CGC genes) and non cancer-associated pc-genes (nonCGC n=19,174). The TSV file containing the CGC data was downloaded from https://cancer.sanger.ac.uk/census with 700 ENSGs with 698 ENSG IDs detected in GENCODE v28 of which 696 are unique (CGC n=696). The same is done for lncRNAs, into CLC2 (n=492) and nonCLC genes (n= 15,314).

#### Matched expression analysis

Based on an in house script used for Survival analysis (section below), TCGA survival expression data for each GENCODE ID is reported and the average FPKM across all tumor samples is calculated. The count distribution of nonCGC and nonCLC gene expression to CGC and CLC2 expression, respectively, is matched using the matchDistribution.pl script (https://github.com/julienlag/matchDistribution).

#### Cancer lncRNA databases

The tested databases were first filtered for lncRNAs in the GENCODE v28 long noncoding annotation (n=15,767).

#### Lnc2cancer

GENCODE IDs from datatable (http://www.bio-bigdata.com/lnc2cancer/download.html) were evaluated (n=512) (40).

#### CRlncRNA

gene names from (http://crlnc.xtbg.ac.cn/download/) were converted to GENCODE IDs (n=146) (38).

#### EVlncRNAs

gene names (http://biophy.dzu.edu.cn/EVLncRNAs/) were converted to GENCODE IDs (n=187) (39).

#### lncRNADisease

gene names from (http://www.rnanut.net/lncrnadisease/index.php/home/info/download) and only cancer-associated transcripts (carcinoma, lymphoma, cancer, leukemia, tumor, glioma, sarcoma, blastoma, astrocytoma, melanoma, meningioma) were extracted. Names were converted to GENCODE IDs (n=137) (37).

### Features of CLC2 genes

#### Genomic classification

The genomic classification was performed as previously (42) using an in house script (https://github.com/gold-lab/shared_scripts/tree/master/lncRNA.annotator).

#### Small RNA analysis

For this analysis “snoRNA”, “snRNA”, “miRNA” and “miscRNA” coordinates were extracted from GENCODE v28 annotation file and intersected with the genomic region of the genes (intronic and exonic regions).

#### Repeat elements

In total 452 CLC2 lncRNAs compared to 1693 expression-matched nonCLC lncRNAs using the LnCompare Categorical analysis (http://www.rnanut.net/lncompare/) (56).

#### Feature analysis

In total 452 CLC2 lncRNAs and 120 mutagenesis lncRNAs are compared to the GENCODE v24 reference using LnCompare (http://www.rnanut.net/lncompare/) (56).

### Cancer characteristic analysis

#### Differential gene expression analysis (DEA)

was performed using TCGA data and TCGAbiolinks. Analysis was performed as reported in manual for matching tumor and normal tissue samples using the HTseq analysis pipeline as described previously. (https://www.bioconductor.org/packages/devel/bioc/vignettes/TCGAbiolinks/inst/doc/analysis.html) (67). For this analysis only matched samples were used and the TCGA data was presorted for tumor tissue samples (TP with 01 in sample name) and solid tissue normal (NT with 11 in sample name). Settings used for DEA analysis: fdr.cut = 0.05, logFC.cut = 1 for DGE output between matched TP and NT samples for 20 cancer types. CLC2 cancer types had to be converted to TCGA cancer types (Supp Fig 6A) Cancer types and number of samples used in the analysis can be found in Supp Fig 6B. DEA enrichment analysis tested with one-sided Fisher’s test. For each CLC2 gene reported as true oncogene (n=275) or tumor suppressor (n=95), hence where no double function is reported (n=22), the positive and negative fold change (FC) values were counted and compared to expression-matched lncRNA genes found in the DEA.

#### Survival analysis

An inhouse script for extracting TCGA survival data was used to generate p values correlating to survival for each gene. Expression and clinical data from 33 cohorts from TCGA with the “TCGAbiolinks” R package (https://bioconductor.org/packages/release/bioc/html/TCGAbiolinks.html) were downloaded (67). P value and Hazard ratio were calculated with the Cox proportional hazards regression model from “Survival” R package (https://cran.r-project.org/web/packages/survival/survival.pdf). All scripts were adapted from here (https://www.biostars.org/p/153013/) and are available upon request. For downstream analysis, only groups with at least 20 patient samples in high or low expression group were used. The plot comprises only the most significant cancer survival p value per gene and was assessed by the Komnogorow-Smirnow-Test (ks-test).

#### Cancer-associated SNP analysis

SNP data linked to tumor/cancer/tumour were extracted from the GWAS page (https://www.ebi.ac.uk/gwas/docs/file-downloads) (n=5,331) and intersected with the whole exon body of the genes. SNPs were intersected to the transcript bed file and plotted per nt in each subset (SNP/nt y axis) and tested using one-sided Fisher’s test.

#### CNV analysis

Human CNV in lncRNAs downloaded from http://bioinfo.ibp.ac.cn/LncVar/download.php (59). NONCODE IDs were converted to GENCODE IDs using NONCODEv5_hg38.lncAndGene.bed.gz. CLC2 and nonCLC ENSGs were matched to NONHSAT IDs with a significant pvalue (0.05, n=733) in the LncVAR table and tested using one-sided Fisher’s test.

#### Code availability

Custom code are available from the corresponding author upon request.

### In vitro validation

#### Cell culture

HeLa and HCT116 were cultured on Dulbecco’s Modified Eagles Medium (DMEM) (Sigma-Aldrich, D5671) supplemented with 10% Fetal Bovine Serum (FBS) (ThermoFisher Scientific, 10500064), 1% L-Glutamine (ThermoFisher Scientific, 25030024), 1% Penicillin-Streptomycin (ThermoFisher Scientific, 15140122). Cells were grown at 37°C and 5% CO2 and passaged every two days at 1:5 dilution.

#### Generation of Cas9 stable cell lines

HeLa cells were infected with lentivirus carrying the Cas9-BFP (blue fluorescent protein) vector (Addgene 52962). HCT116 were transfected with the same vector using Lipofectamine 2000 (ThermoFisher Scientific, 11668019). Both cell types were selected with blasticidin (4ug/ml) for at least five days and selected for BFP-positive cells twice by fluorescence activated cell sorting.

#### CRISPR inhibition sgRNA pair design and cloning

sgRNA pairs targeting *LINC00570* were designed using GPP sgRNA designer (https://portals.broadinstitute.org/gpp/). The sgRNA pairs were manually selected from the output list and cloned into the pGECKO backbone (CRISPRi.1: *5’ GTTACTTCCAACGTACCATG 3’*, CRISPRi.2: *5’ CCTGTACCCCCATGGTACGT 3’*) (Addgene 78534; (68))

#### Antisense LNA GapmeR design

Antisense LNA GapmeR Control (*5’ AACACGTCTATACGC 3’)* and three Antisense LNA GapmeR Standard targeting *LINC00570* (LNA1: *5’ GGAAATTGCTCTGATG 3’*, LNA2: *5’ GATTGGCATTGGGATA 3’*, LNA3: *5’ GAAGTGGCCTGAGAAA 3’*) were designed and purchased at Qiagen.

#### RT-qPCR

For each time point total RNA was extracted and reverse transcribed (Promega). Transcript levels of *LINC00570* (FP: *5’ TAGGAGTGCTGGAGACTGAG 3’*, RP: *5’ GTCGCCATCTTGGTTGTCTG 3’*) and housekeeping gene *HPRT1* (FP: *5’ ATGACCAGTCAACAGGGGACAT 3’*, RP: *5’ CAACACTTCGTGGGGTCCTTTTCA 3’*) were measured using GoTaq qPCR Master Mix (Promega, A6001) on a TaqMan Viia 7 Real-Time PCR System. Data were normalized using the ΔΔCt method (69)).

#### Gene-specific RT-PCR and cDNA amplification

From the extracted total RNA, we performed a gene specific reverse transcription using the reverse primers for *LINC00570* and *HPRT1* to enrich for their cDNA. Presence or absence of transcript was detected by a regular PCR using GoTaq^®^ G2 DNA Polymerase (Promega, M7841) from 100ng cDNA and visualized on an agarose gel.

#### Viability assay

HeLa and HCT116 cells were transfected with Antisense LNA GapmeRs at a concentration of 50nM using Lipofectamine 2000 (Thermofisher) according to manufacturer's protocol. One day after, transfected cells were plated in a white, flat 96-well plate (3000 cells/well). Viability was measured in technical replicates using CellTiter-Glo 2D Kit (Promega) following manufacturer's recommendations at 0, 24, 48, 72 hours after seeding. Luminescence was detected with Tecan Reader Infinite 200. Statistical significance calculated by t-test.

For CRISPR inhibition experiments, HeLa-Cas9 and HCT116-Cas9 cells were transfected with control sgRNA plasmid and two *LINC00570* targeting plasmids. Cells were selected with puromycin (2ug/ml) for 48h. Viability assay was performed as previously described.

## Supplementary Data

**Supplementary Table 1:** Excel table with all CLC2 and cancertype and evidence level

**Supplementary Table 2:** Excel table with all CLC2 ENSG with cancer functionality

## Supplementary Figures

**SUPP. Figure 1:**
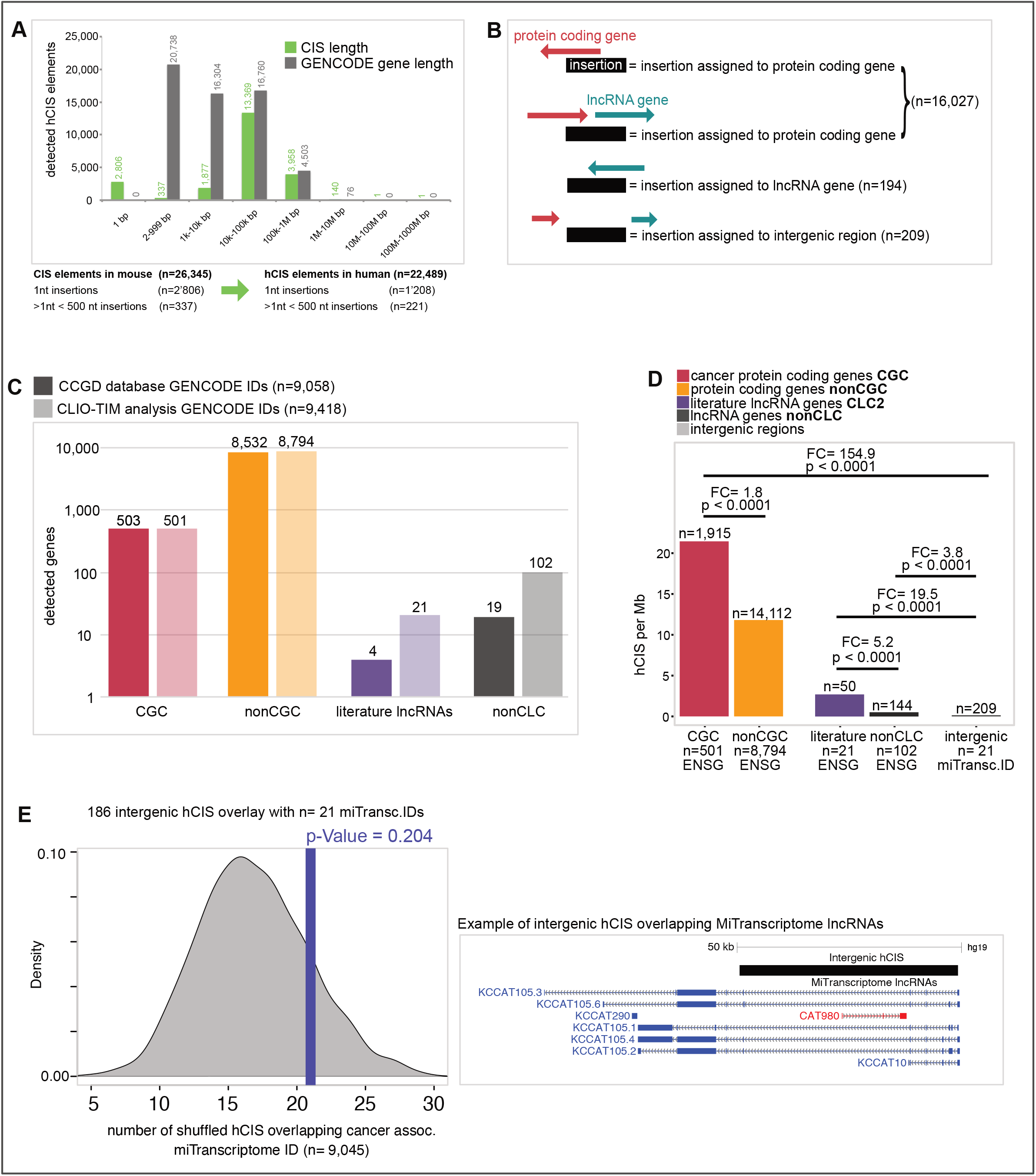
Insertion analysis **A)** All insertion sizes after Liftover compared to GENCODE v28 gene length. Number of input CIS elements in mouse compared to hCIS elements in human after Liftover. **B)** Assign genes to GENCODE v28 genes and gene families. C) CCGD reported genes (dark) and CLIO-TIM reported genes (fade) for each gene class. **D)** Genes with insertion categorized in gene types. Statistical significance calculated by one-sided Fisher’s test. **E)** Assign intergenic regions to MiTranscriptome IDs and compare to shuffled hCIS overlayed with MiTranscriptome IDs. Example of one insertion site with MiTranscriptome ID.

**SUPP. Figure 2:**
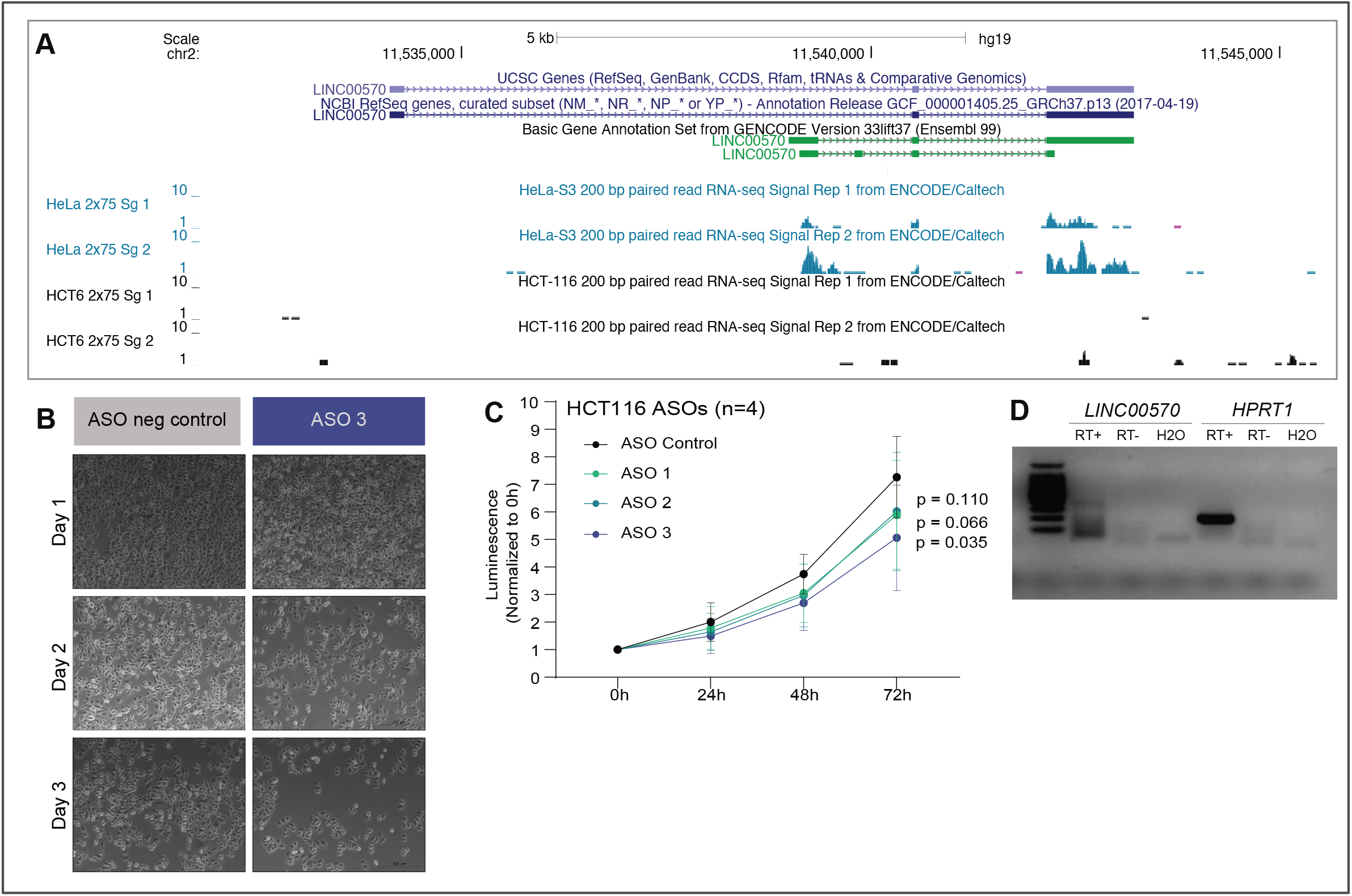
*LINC00570* insertion candidate characteristics. **A)** ENCODE expression data of *LINC00570* in HeLa (blue) and HCT116 cells (black). **B)** Cell proliferation of HeLa cells treated with ASO negative control and ASO 3 at Day 1, 2 and 3. **C)** Proliferation assay for HCT116 cells with ASO control and the three ASO targeting *LINC00570*. Statistical significance calculated by Student’s *t*-test. **D)** RT-PCR of *LINC00570* and *HPRT1* from HCT116 cells to check for expression.

**SUPP. Figure 3:**
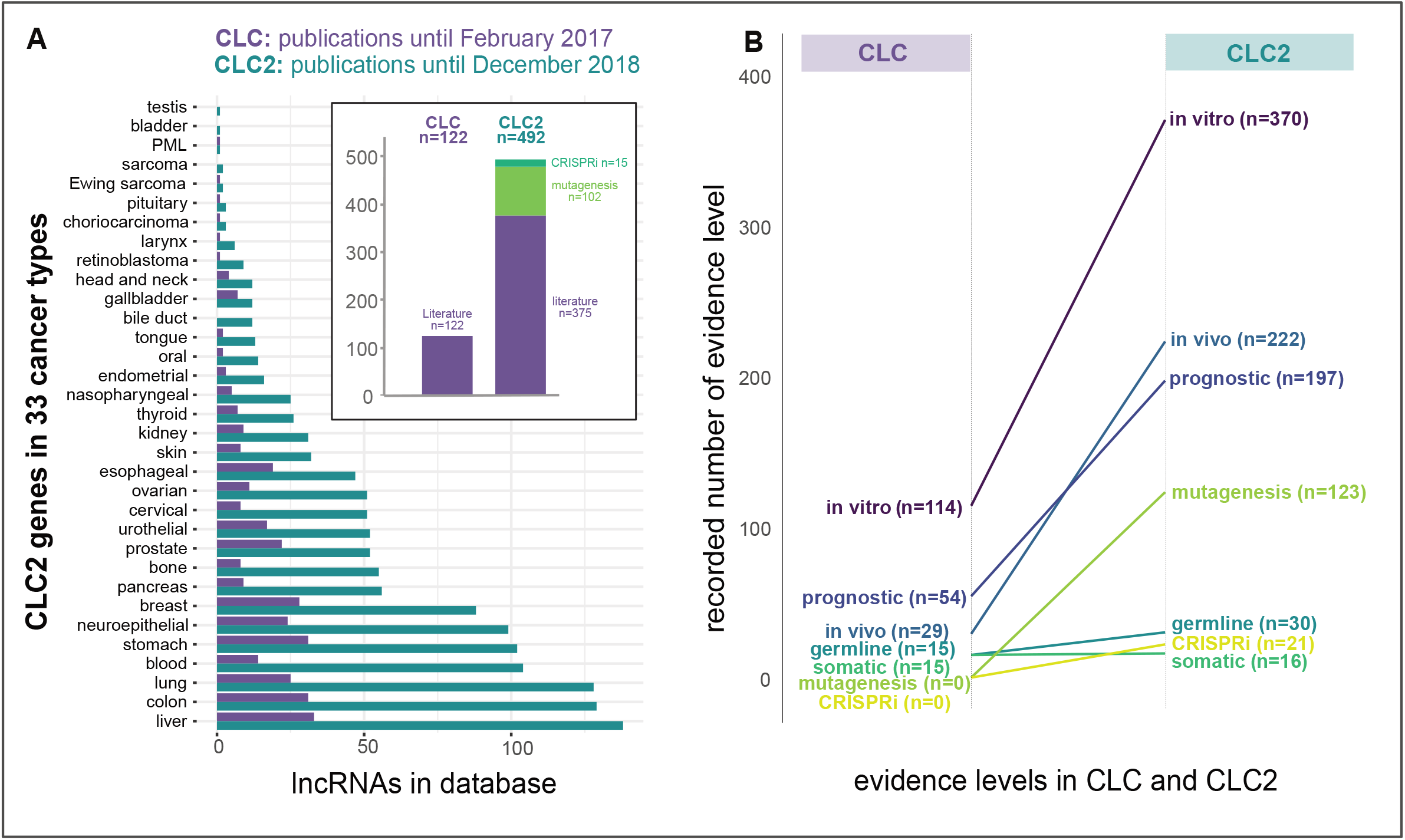
comparison of first CLC and CLC2. **A)** CLC2 genes are detected in 33 cancer types and compared to 29 in the first CLC. CLC reported n=122 literature lncRNAs whereas CLC2 comprises 492 genes from 3 different analysis. **B)** Comparing evidence levels of genes from the initial CLC with the CLC2.

**SUPP. Figure 4:**
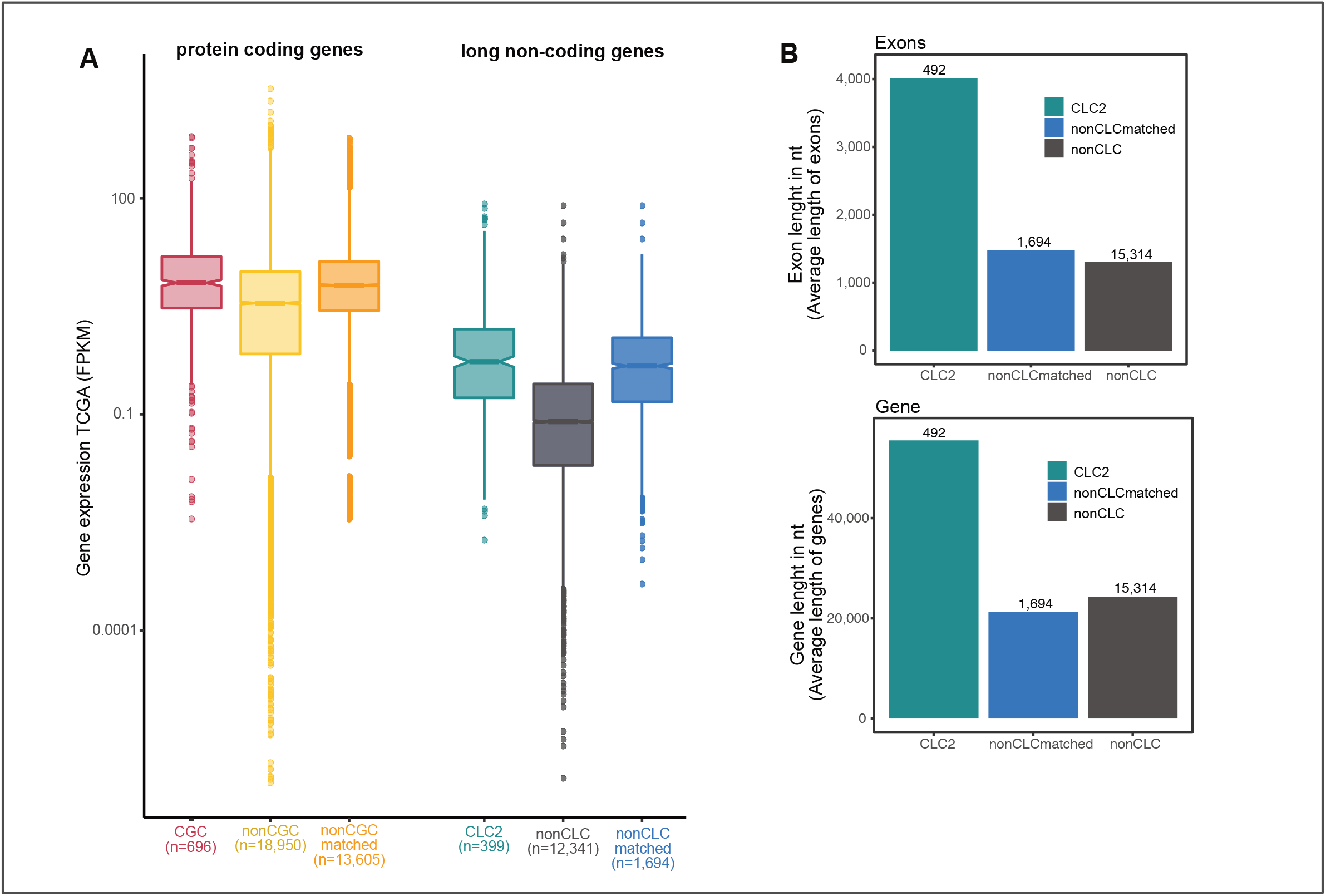
CLC2 expression and gene length bias. **A)** CLC2 genes (turquoise) are higher expressed than nonCLC genes (grey), same for CGC genes (red) compared to nonCGC genes (orange). Expression-matched CLC2 (blue) and CGC (yellow) were generated and match the expression of the CLC2 and CGC, respectively. **B)** CLC2 genes (turquoise) with increased exon and whole gene body length when compared to expression-matched (blue) and all other lncRNAs (grey).

**SUPP. Figure 5:**
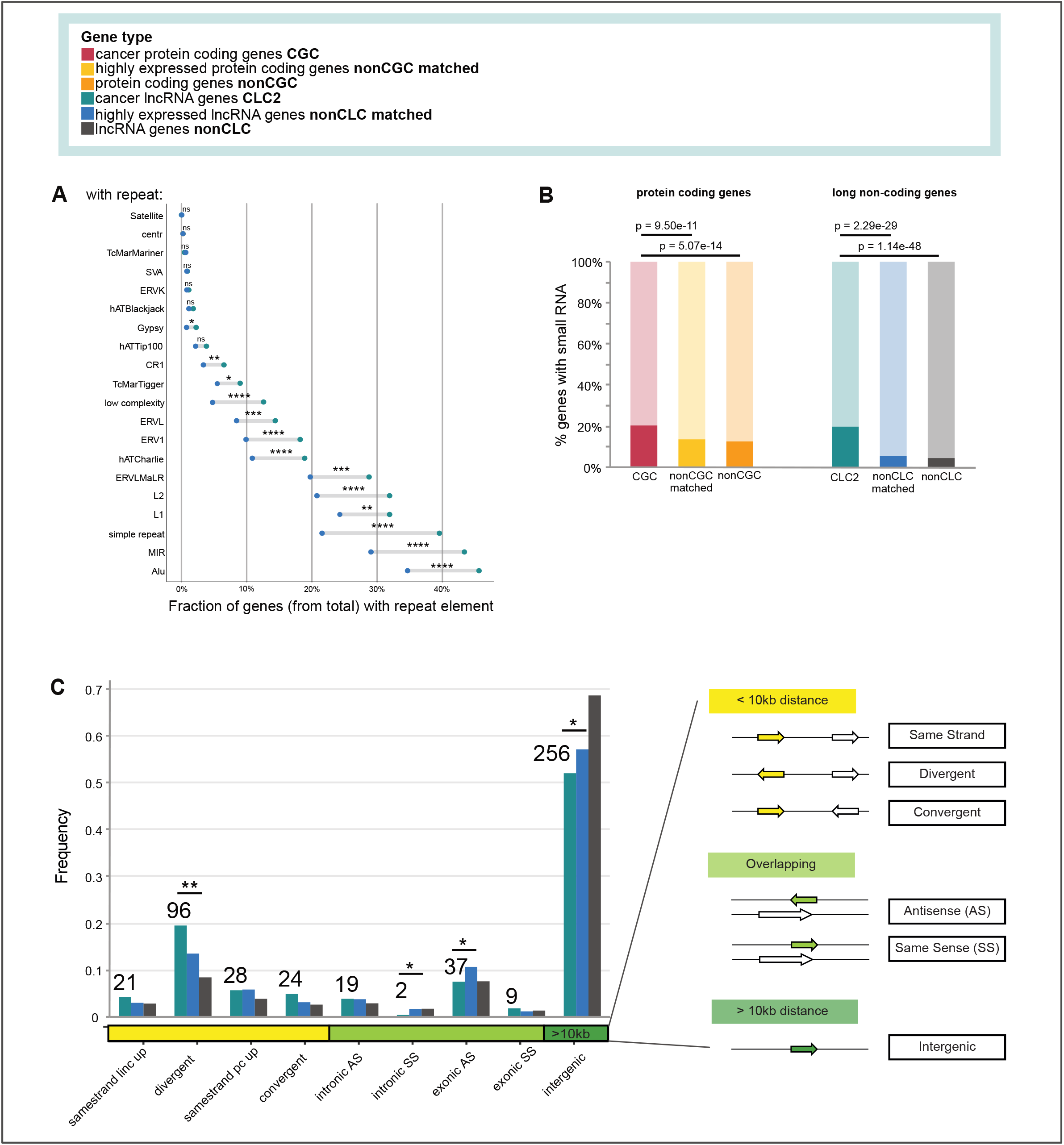
CLC2 gene characteristics. **A)** CLC2 genes are enriched for ⅔ of the analyzed repeat element families when compared to expression-matched nonCLC genes. Statistical significance calculated by hypergeometric test (highly significant ****=<0.0001). **B)** CGC and CLC2 genes are enriched for small RNAs compared to expression-matched nonCGC and nonCLC, respectively. In the bar graph we report the fraction of genes of each dataset with (dark color) or without (light color) small RNA encoded in the genomic region. Statistical significance calculated by one-sided Fisher’s test. **C)** Genomic classification of CLC2, expression-matched nonCLC and nonCLC genes. Statistical significance calculated by two-sided Fisher’s test (*=<0.05).

**SUPP. Figure 6:**
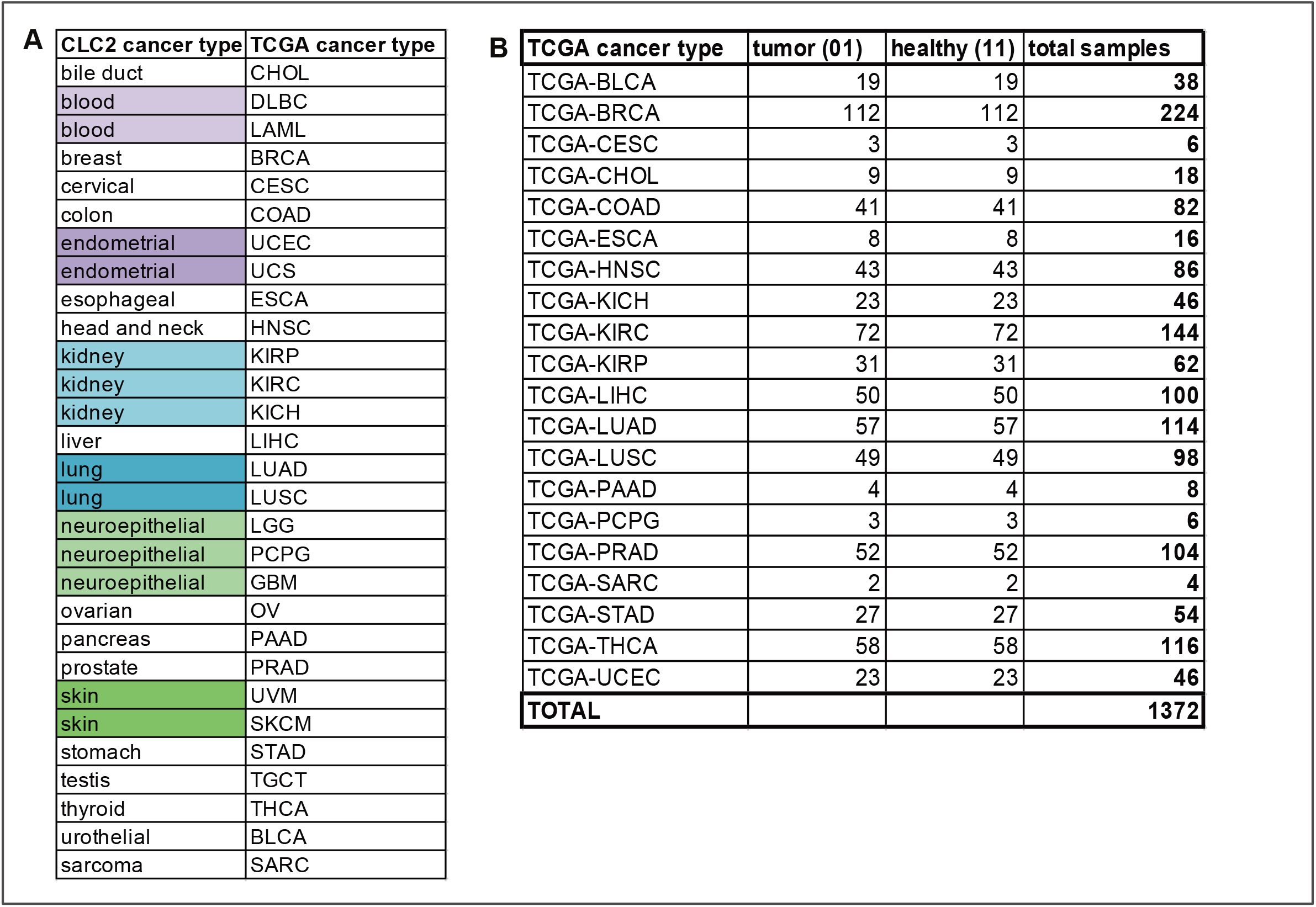
TCGA cancer types for differential expression analysis. **A)** CLC2 cancer types corresponding to TCGA cancer types. **B)** Samples for each TCGA cancer type analyzed for differential expression analysis.

**SUPP. Figure 7:**
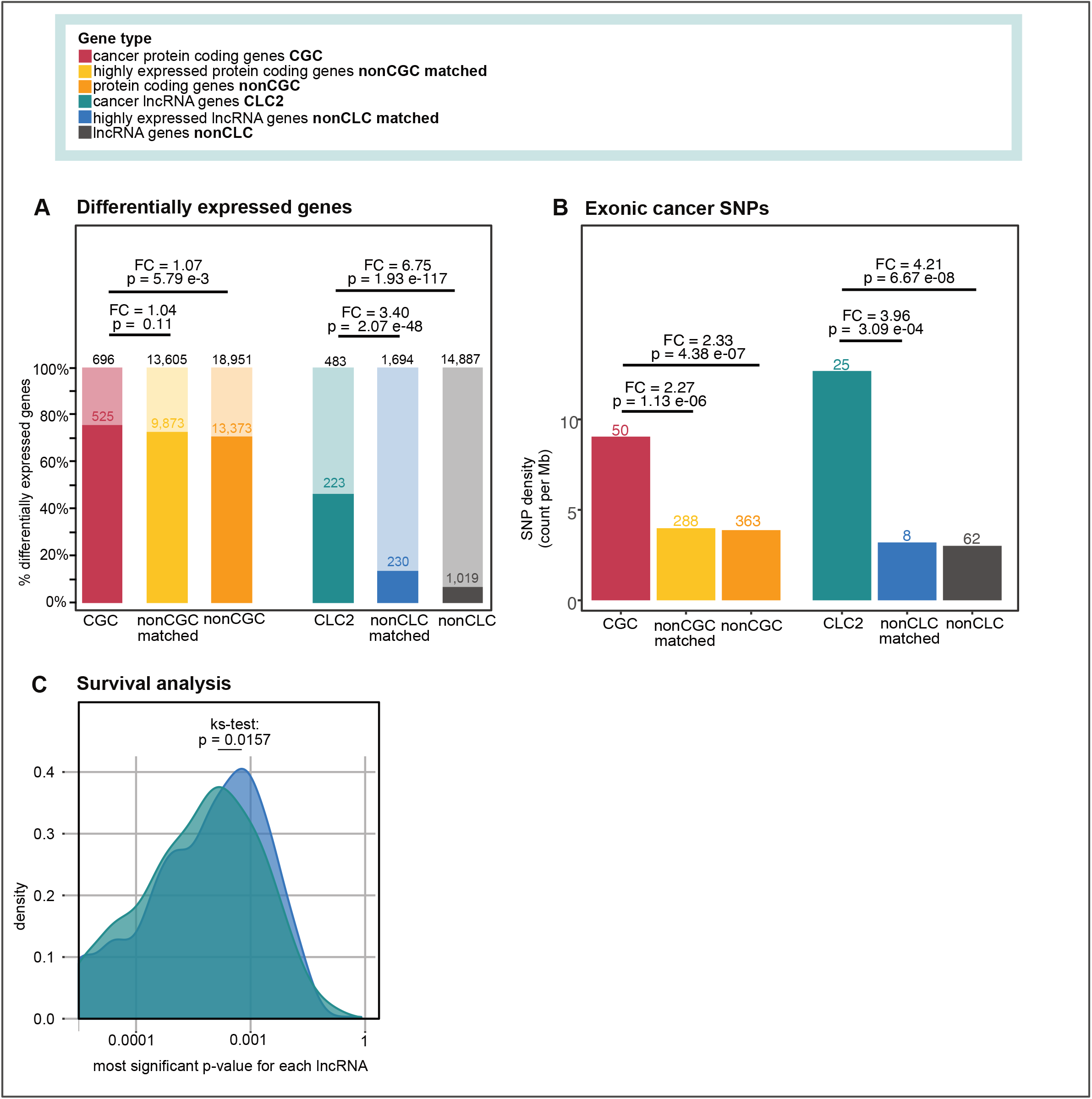
Cancer characteristics for all analyzed gene types. **A)** Differentially expressed genes enriched in cancer-associated gene families (CGC and CLC2). Statistical significance calculated by one-sided Fisher’s test. **B)** exonic cancer SNPs enriched in cancer-associated gene families (CGC and CLC2). Statistical significance calculated by one-sided Fisher’s test. **C)** survival analysis comparing most significant p-value for each lncRNA in the CLC2 compared to expression-matched lncRNAs. Statistical significance calculated by ks-test.

## Acknowledgements

We gratefully acknowledge administrative support from Ana Radovanovic and Silvia Roesselet (DBMR, University of Bern). We also acknowledge Joana Carlevaro-Fita (EPFL, Lausanne) and Judith Bergada (University of Zurich) for the helpful advice and discussions as well as Roberta Esposito, Panagiotis Chouvardas, Hugo Guillen Ramirez and the other members of the Laboratory for Genomics of LncRNA and Disease for their valuable input.

## Author contribution

RJ conceived the project. RJ, AV, AH performed manual annotation of CLC2. AV performed the feature analysis, evolutionary analysis, mutation analysis, differential expression, GWAS SNP, CNV analysis and data integration. AL performed survival analysis. NB performed the ASO and CRISPRi KD experiments. AV, NB and MT performed the qPCR experiments. RJ, AV, AL, NB, MT and SH drafted the manuscript and prepared the figures and supplementary material. All authors read and approved the final draft.

## Conflict of interest

The authors declare that they have no competing interests.

## The Paper Explained

Problem: Cancer is one of the leading causes of death worldwide. The development of effective therapies depends on creating collections of known cancer genes. These can comprise not only conventional protein coding genes, but also more recently discovered genes like long noncoding RNAs (lncRNAs). LncRNAs are considered highly promising therapeutic targets, however the relatively poor state of knowledge, and the lack of high quality cancer lncRNA collections, represents a significant hurdle to developing lncRNA therapies.

Results: To address the need for collections of cancer lncRNAs, we have developed the Cancer lncRNA Census 2 (CLC2). CLC2 consists of 492 cancer lncRNAs functionally validated in 33 cancer subtypes. CLC2 is the first catalogue to incorporate automatic screen data from mice, and is shown to be superior to existing collections across several criteria. We show that CLC2 lncRNAs enriched for cancer associated mutations and tend to be differentially expressed in tumours. Impact: CLC2 is a critical resource for future development of cancer therapies targeting lncRNAs. Analysis of these genes has provided new insights into their biological and clinical properties.

## Ethics approval and consent to participate

Not applicable.

## Consent for publication

Not applicable.

## Availability of data and materials

Information on CIS elements for mouse and human lncRNAs reported in this publication are available in the Supplementary Table 1 and the code is available from the corresponding author on request.

## Funding

This work was funded by the Swiss National Science Foundation through the National Center of Competence in Research (NCCR) “RNA & Disease”, by the Medical Faculty of the University and University Hospital of Bern, by the Helmut Horten Stiftung and Krebsliga Schweiz (4534-08-2018).

## Notes

### Competing Interest Statement

The authors have declared no competing interest.

